# Measurement of cellular traction forces during confined migration

**DOI:** 10.1101/2024.09.27.615466

**Authors:** Max A. Hockenberry, Andrew J. Ulmer, Johann L. Rapp, Frank A. Leibfarth, James E. Bear, Wesley R. Legant

## Abstract

To migrate efficiently through tissues, cells must transit through small constrictions within the extracellular matrix. However, in vivo environments are geometrically, mechanically, and chemically complex, and it has been difficult to understand how each of these parameters contribute to the propulsive strategy utilized by cells in these diverse settings. To address this, we employed a sacrificial micromolding approach to generate polymer substrates with tunable stiffness, controlled adhesivity, and user-defined microscale geometries. We combined this together with live-cell imaging and three-dimensional traction force microscopy (TFM) to quantify the forces that cells use to transit through constricting channels. Surprisingly, we observe that cells migrating through compliant constrictions take longer to transit and experience greater nuclear deformation than those migrating through more rigid constrictions. TFM reveals that this deformation is generated by inwardly directed contractile forces that decrease the size of the opening and pull the walls closed around the nucleus. These findings show that nuclear deformation during confined migration can be accomplished by internal cytoskeletal machinery rather than by reactive forces from the substrate, and our approach provides a mechanism to test between different models for how cells translocate their nucleus through narrow constrictions. The methods, analysis, and results presented here will be useful to understand how cells choose between propulsive strategies in different physical environments.

**Significance Statement:** Cell migration is critical for both physiological events like wound healing and pathological events like metastasis. Understanding how cells move through complex environments will assist efforts to enhance or inhibit such processes. We developed a method to quantify the forces that cells use to move through multidimensional environments, including through narrow constrictions like those in tissues. Surprisingly, we find that cells transiting through soft constrictions take longer and deform more than those transiting through rigid constrictions, and we connect this finding to inwardly directed contractile forces generated by migrating cells. Together, this work reveals a key role for substrate rigidity to regulate cell transit through confining geometries and provides a quantitative platform to investigate similar processes in other settings.

## Introduction

Cells migrating *in vivo* encounter varying geometric and mechanical environments. Often these environments contain spatial constrictions that are smaller than the size of individual cells. To effectively transit through such constrictions, cells must deform their nucleus, which is the largest and most rigid organelle (1). Previous studies have identified that smaller constrictions require greater nuclear deformation and can restrict cell migration (2, 3). It has been suggested that cells can push (4, 5), pull (6–8), and actively deform the nucleus using internal cytoskeletal machinery (3, 4, 9) to pass through small constrictions. However, it is less defined how the mechanical properties of the constriction and surrounding region might regulate confined migration. Additionally, it is unclear how and where cells exert the forces needed to physically propel themselves through a constriction and how substrate rigidity and geometry might influence this process. Without direct measurement of the forces between the cell and surrounding matrix, it has been challenging to distinguish between these physical mechanisms.

Traction force microscopy (TFM) is a powerful method for measuring the forces that cells use to migrate in both 2D and 3D settings (10–12). This approach has revealed that cells use actomyosin machinery to generate contractile forces which are exerted through sites of substrate adhesion (13–15). These forces induce deformations in the substrate that can be measured by imaging the motion of fiducial markers relative to a cell-free reference state. The deformations are then converted to tractions using *a-priori* knowledge of the substrate mechanics and geometry (10). TFM has been an essential tool to understand the relationship between traction forces and motility (16– 18) and to dissect how cells physically navigate through their environment. Previous work has established 3D TFM pipelines for cells embedded within homogeneous hydrogels (11), crawling within grooves or through narrow tubes formed from polyacrylamide (19, 20), or on poly (dimethyl siloxane) (PDMS) micropillars in confining microdevices (21). These systems have shown that cells on 2D surfaces (14), within grooved channels (20), and within 3D hydrogels (11) all exert inwardly directed contractile forces. In contrast, initial reports for cell-generated forces in confining 3D channels (19) or between two confining planar surfaces (22) have reported both inward and outward directed cell forces depending on cell type and adhesive conditions. Together, these studies demonstrate the role of substrate geometry in regulating cellular forces. However, it remains challenging to precisely pattern substrate geometry at micron scales, to control substrate rigidity over physiological ranges, and to measure forces with sub-cellular resolution which makes it challenging to integrate the results across varying experimental conditions. Overcoming these challenges would allow for testing of the complex relationship between substrate geometry, substrate rigidity, and the migration strategies that cells adopt to navigate confinements in various biologically relevant contexts.

In this work, we develop a sacrificial micromolding technique to generate PDMS substrates with micron scale geometric features and physiologically relevant stiffnesses. PDMS substrates do not swell under common experimental conditions which makes it possible to reproducibly pattern microscale features such as confinements (23). We combine this with 3D TFM and live-cell microscopy to measure the forces generated by cells migrating within user-defined 3D channels with small constrictions. Surprisingly, we find that cells migrating through compliant constrictions had greater nuclear deformation and impaired transit compared to those in rigid constrictions. 3D TFM revealed that this phenomenon is caused by inwardly directed contractile forces that pull the channel walls closed around the cell, even within the constricting region. Based on this, we conclude that, at least under the geometries and cell types studied here, the nuclear deformation necessary to transit confinements is produced via internal cytoskeletal machinery rather than reaction forces from pulling or pushing the nucleus through the constriction.

## Results

### Generation of compliant microchannels with user-defined micron-scale features

Due to the need for mechanical demolding, it is difficult to generate compliant substrates with micron scale features using standard soft lithography approaches (24). To overcome this, we adapted a sacrificial micro-molding approach (25). We developed a workflow with three micropatterning steps. First, we used photolithography to create raised micropattern features in photoresist on silicon wafers and cast a negative replica of the photoresist pattern using rigid PDMS (**Figure 1A (i,ii)**). While this mold faithfully transforms the raised photoresist features into recessed PDMS channels, the resulting substrate is substantially more rigid than most biological tissues and cannot be used to measure cell-generated forces. Next, we recast this PDMS mold with a sacrificial material made from molten sugar (**Figure 1A (iii**,**iv)**). Lastly, we cast the sugar template with a soft PDMS mixture (of Sylgard 527 and Sylgard 184) and allowed it to fully cure before dissolving the sugar in water (**Figure 1A (v,vi)**). This method avoids the need to physically demold the soft PDMS substrate and maintains high-fidelity features (**Figure 1A**). We can tune the rigidity of the final substrate over a range of physiologically relevant values by varying the ratio of Sylgard 527 and Sylgard 184, (**Figure 1B, Supplemental Figure 1A-C**) and fabricate a variety of surface features by controlling the design of the mask used to pattern the original photoresist template (**Figure 1C, Supplemental Figure 1D**). Importantly, in contrast to polyacrylamide or polyethylene glycol, which are commonly used for TFM (23), the final PDMS template is a non-hydrogel elastomer that is dimensionally stable under changes to temperature, osmolarity, and aqueous solvents.

**Figure 1.**
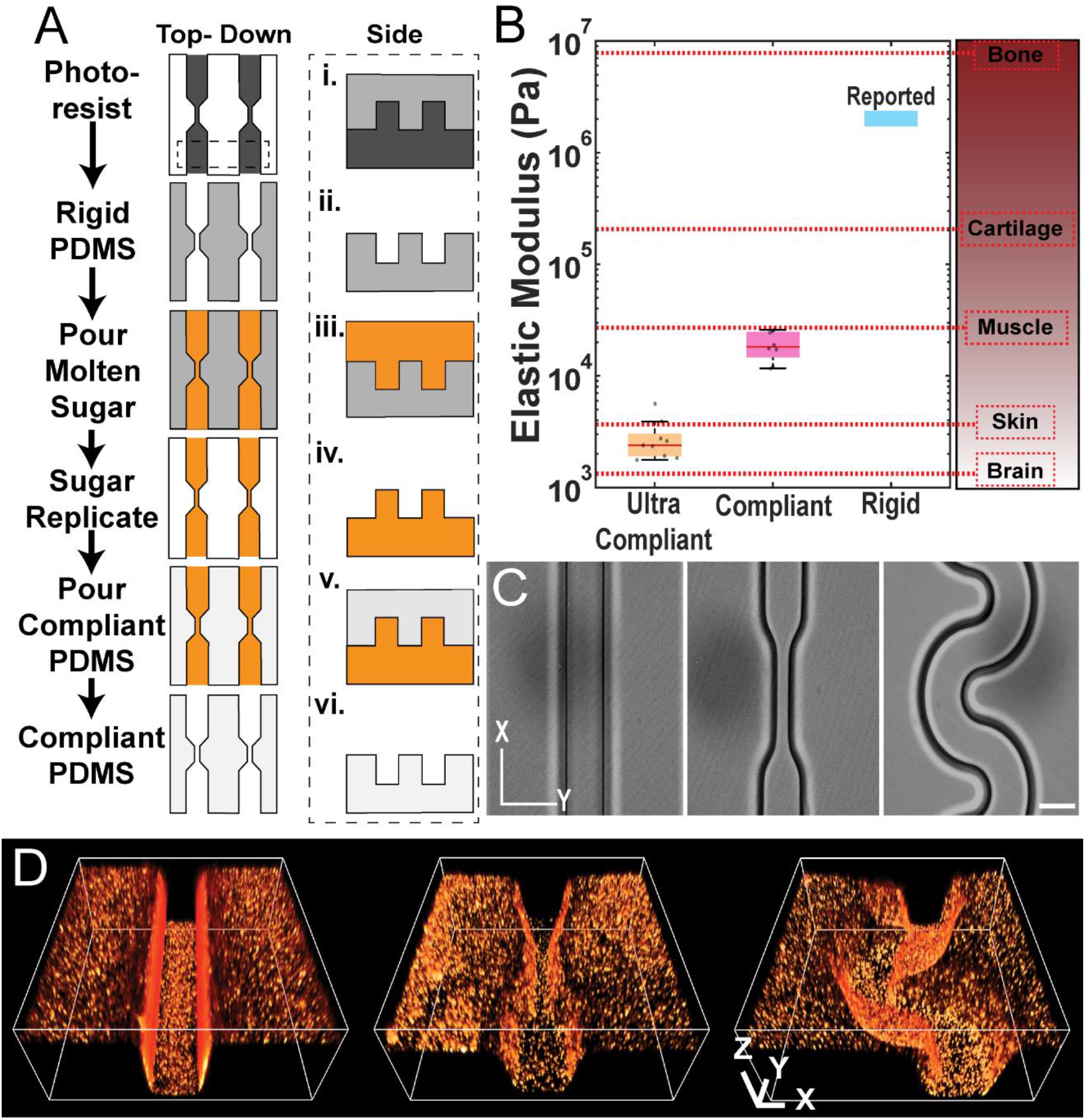
Fabrication of user-defined compliant PDMS substrates. **A)** Silicon wafers of desired patterns generated with standard photolithography (dark gray), rigid Sylgard 184 PDMS (medium gray), and sacrificial sugar micromolds (orange) are used to generate compliant substrates (light gray) of user-defined patterns. **B)** Measured elastic moduli of different PDMS formulations and relative to that of different tissues. **C)** Brightfield images of straight channels, confinements, and radial curvatures in compliant PDMS. Scale bar = 20 microns. **D)** Volume renderings of PDMS microstructures coated with fluorescent microspheres. Bounding boxes = 100×100 × 44 µm.

Furthermore, it is amenable to covalent addition of fluorescent beads (**Figure 1D**) which allows for tracking of cell induced displacements during migration and functional coating of extracellular matrix (ECM) ligands such as fibronectin (**Supplementary Figure 1E**) to control cell adhesion.

### Imaging of confined cell migration in rigid and compliant microchannels

We first sought to test the effect of substrate rigidity on the ability of cells to transit through a constriction. To do so, we cultured mouse embryo fibroblast (MEF) cells in substrates consisting of thirty-micron deep channels with forty micron long by five-micron wide constrictions every 200 microns along the channel (**Fig 2A, B**). This geometry closely matches similar geometries used in other studies, and the constriction is of a similar size to those encountered in vivo (2–6, 9, 26–28). To study the effects of substrate rigidity, we chose to compare 20 kPa compliant (similar to muscle tissue (29)) to the rigid PDMS (approximately 2 MPa) most commonly used in previously reported confined migration assays. We then captured transmitted light and fluorescence microscopy images of migrating cells and their nuclei under low (10X) magnification. A large field of view enabled us to capture dozens of events of cells transiting through the channel confinements in a single 24-hour experiment (**Figure 2A, B, Supplemental Movie 1**). This provided sufficient statistical power to detect clear differences in cells transiting through compliant vs. rigid constrictions (**Figure 2A, B**). Contrary to our expectations, cells on compliant substrates were significantly slower to transit constrictions compared to those on rigid substrates (**Figure 2C**). Closer examination of the images suggested that during transit of compliant constrictions, the cells pulled the channel walls inward, effectively decreasing the open width of the constriction, which we confirmed by measuring the change in channel width from relaxed to maximum deformation (**Figure 2D, Supplemental Movie 2**). Additionally, we observed the nuclei of cells transiting through compliant constrictions were more deformed and had a higher aspect ratio than those in rigid constrictions (**Figure 2E, 2F**).

**Figure 2.**
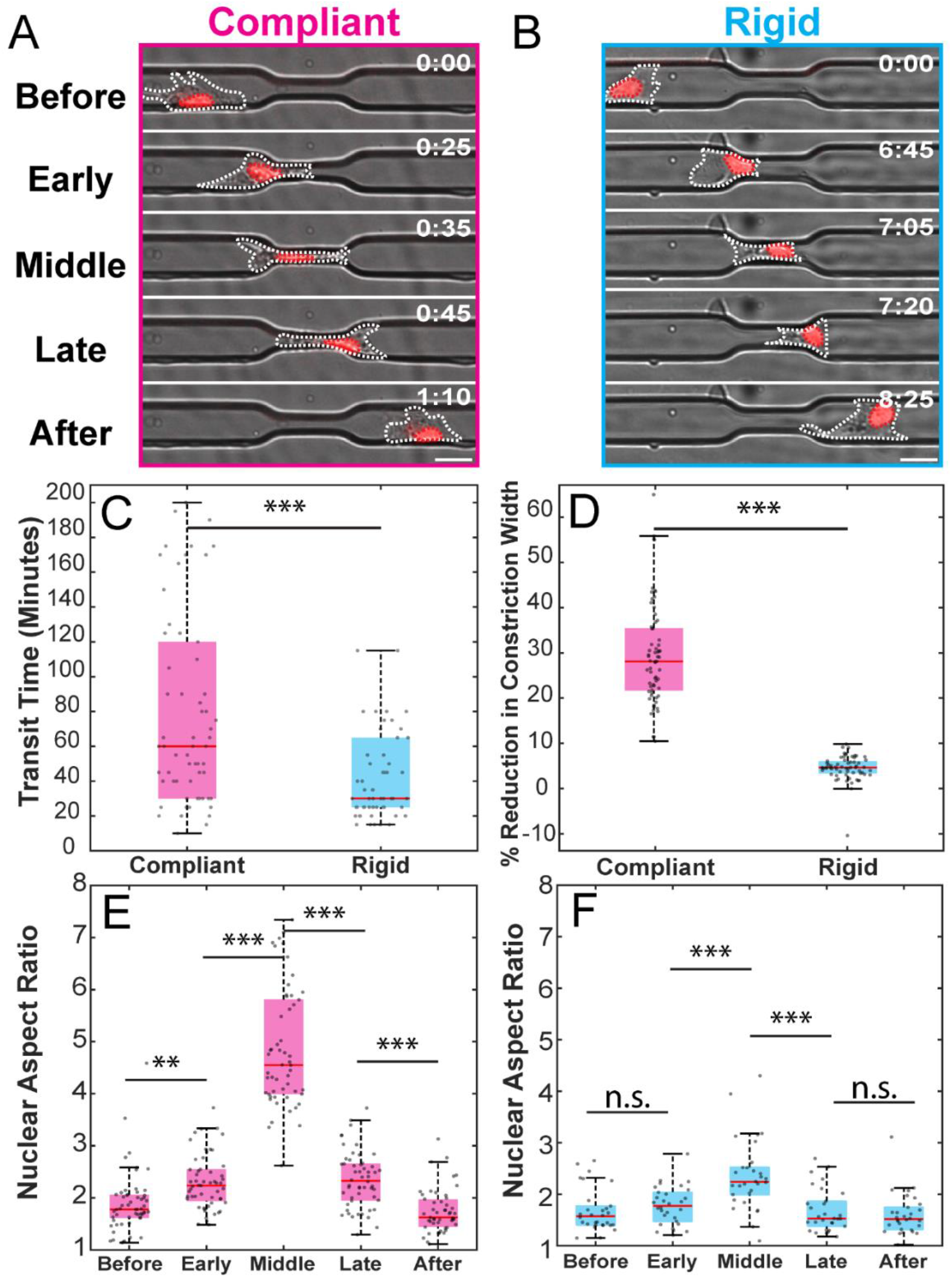
Cells undergoing confined migration in compliant substrates pull channel walls towards themselves. **A, B)** Montage images of cells passing through 5-micron wide confinements in compliant (A) and rigid (B) microchannels. Time is hours:minutes. **C)** Transit time required for the centroid of the nucleus to pass through the 40-micron long, 5-micron wide confinements. **D)** Maximum percent reduction in the constriction width on compliant and rigid substrates during nuclear transit. **E, F)** Nuclear aspect ratio during confined migration at distinct stages of nuclear transit in compliant (E) and rigid (F) substrates. *** is p<0.001, ** is p<0.01, and * is p<0.05 as determined by Students T Test (**Panels D, E**, and **F**, Bonferroni correction for multiple comparisons applied for Panel E and F) while **Panel C** uses the Wilcoxon rank-sum test. N > 50 cells from at least 3 independent biological replicates.

### Measurement of deformations exerted by cells in compliant microchannels

Although we expected that transiting cells would cause greater displacements in compliant rather than in rigid constrictions, we were surprised by the direction of these displacements. Instead of the walls of the complaint constriction being pushed outward by the transiting nucleus, we observed that the walls were pulled inward by the cell, which resulted in greater nuclear deformation than was observed in rigid channels. To better understand how cells generated these deformations, we sought to measure the cell-generated forces as they transited through constrictions. To do so, we adapted finite-element-based TFM to explicitly incorporate the micropatterned geometry of the substrate ((14), **Methods**). We quantitatively mapped the substrate displacements by coating the surface with 100 nm fluorescent beads and capturing 3D volumetric timelapse movies using spinning disc confocal microscopy (**Figure 3A - C**). At the end of the experiment, we removed cells using the detergent SDS and acquired a final reference image of the non-deformed substrate. We localized the bead centroids in both the timelapse and the reference images and then computed the displacement field using previously described methods (11, 14). With this approach, we observed inwardly directed displacements at the leading and trailing edges of the cell. Qualitatively, the leading edge was frequently “Y” shaped and spanned the channel walls while the trailing edge displayed a single point of attachment. Together, these patterns suggested that the cells often generated contractile tripoles that are oriented along the channel axis (**Figure 3B**).

**Figure 3.**
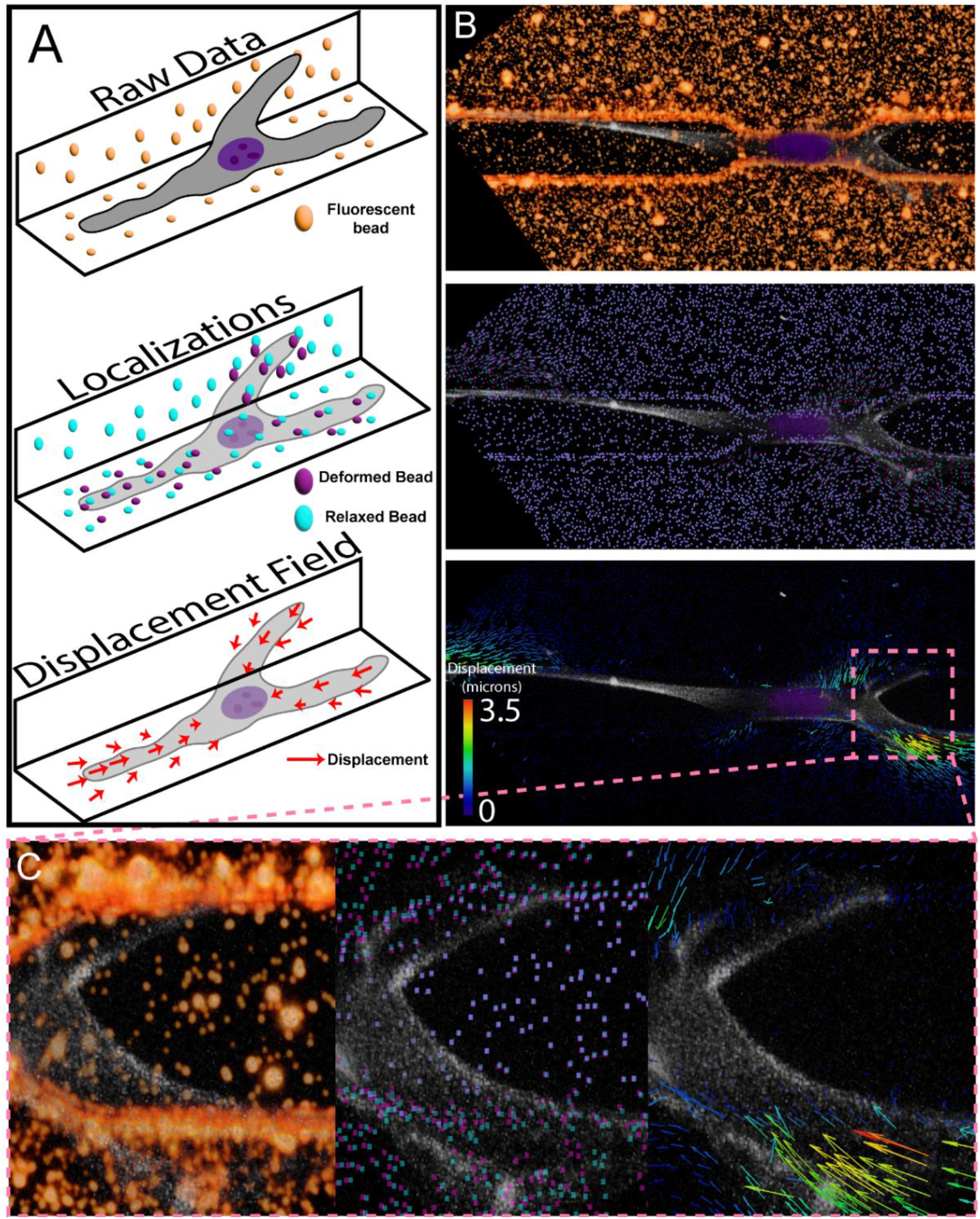
Calculation of 3D displacement fields on compliant substrates with user defined geometries. **A)** Cartoon diagram of displacement field calculation. Raw images of fluorescent beads are converted into displacement fields by first localizing beads in the reference (cell free, cyan) state, and then localizing beads in the experimental frame (purple). Beads are then linked by a feature vector algorithm. **B)** Representative volumetric data set showing raw image data (top), localized bead positions (middle), and resulting displacement field (bottom) as overlaid vectors. Vector lengths in the bottom panel are exaggerated for visualization. **C)** Zoomed in view of a high bead displacement region showing the raw image data (left), localized bead positions (center), and resulting displacements (right) as overlaid vectors. Vector lengths in the right panel are exaggerated for visualization.

### Quantification of 3D cellular traction forces during migration in compliant microchannels

To calculate the cell-generated traction forces from 3D displacement fields, we utilized finite element modeling to generate a geometry-specific Green’s matrix as described previously ((11, 14), **Figure 4A-E**). In brief, we utilized the 3D fluorescent bead coordinates from the reference (non-cell containing) volume to generate a 3D tetrahedral mesh of the substrate geometry (**Figure 4A**). Using this mesh, we computed the predicted displacement in response to simulated unit tractions applied in each of the Cartesian directions at each facet on the surface. (**Figure 4B, C**).

**Figure 4.**
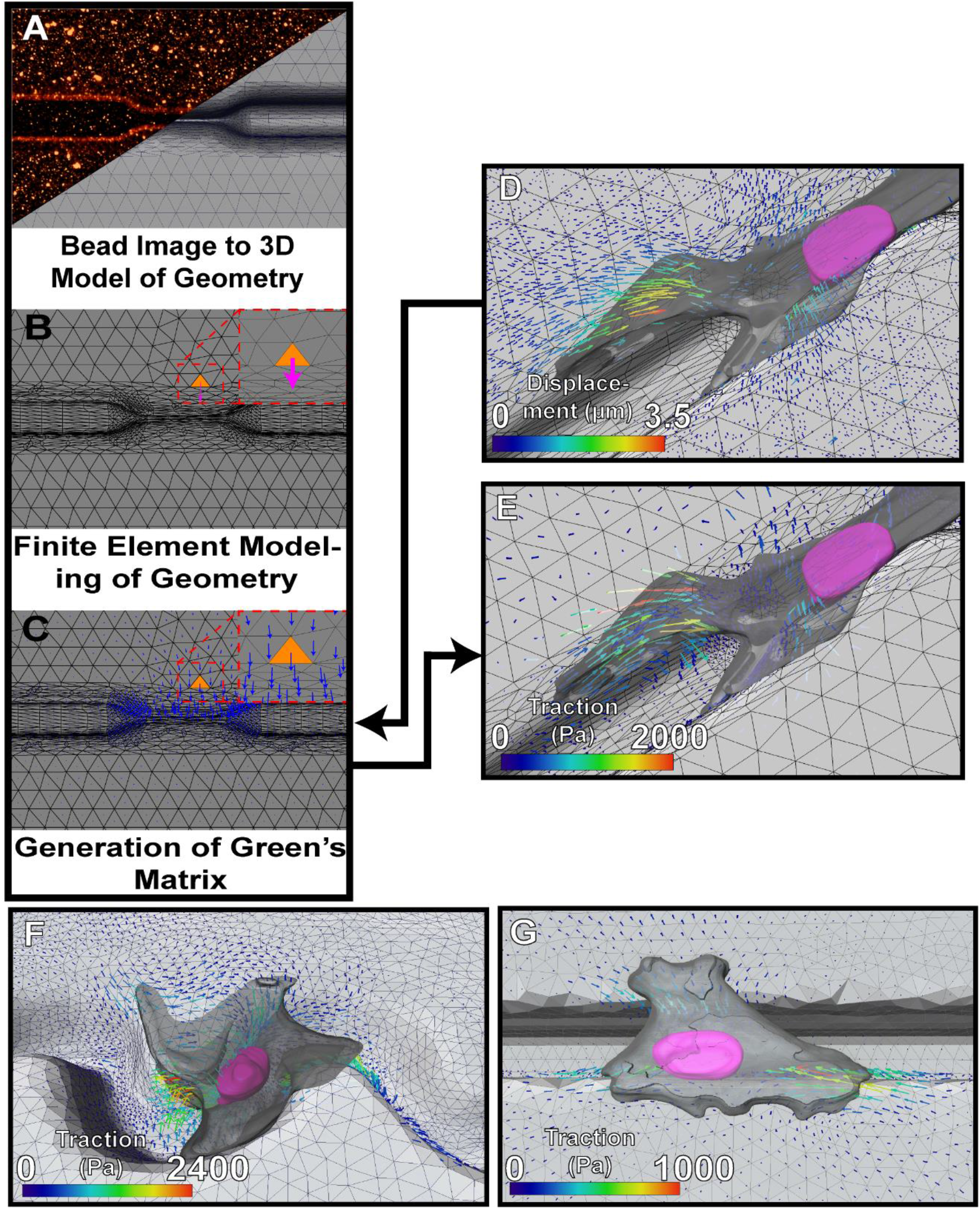
Calculation of 3D traction fields on compliant substrates with user defined geometries. **A-C)** Flow chart for the calculation of traction forces. 3D images of the fluorescent beads are used to generate a 3D mesh of the microchannel geometry (A). This mesh is then used for finite element modeling where a unit load is applied to each element of the mesh surface (B) which results in a Green’s matrix to convert a displacement field to a traction field (C). **D)** Experimentally measured displacement fields are decomposed by inverting the Green’s matrix to compute traction fields (**E**). **E-G)** Representative traction fields for cells in a confinement (E), radial curvature (F) and straight channel (G).

Together, these finite element solutions can be assembled into a Greens’ matrix for the system that describes how forces on the surface map to displacements of the substrate. This can then be used to compute the traction field from a given displacement field (**Figure 4D - G, Supplementary Movie 3**).

### Geometric constraint in migrating cells induces polarization, increased strain energy, and anisotropic traction forces

Previous studies have suggested that the cell nucleus represents a major impediment to constricted migration. While the cytoplasm is relatively fluid and deformable, the nucleus is substantially stiffer (1) and the cell must generate sufficient force to deform and propel it through a constricting region. To better understand how cells migrate on compliant substrates and generate the forces necessary to enable transit in confinements, we visualized migrating MEFs labeled with a live-cell compatible DNA dye. We plated these cells in compliant channels and measured the forces that cells generated while migrating through 5-micron constrictions as compared to cells migrating through channels without confinements as a control (**Supplemental Figure 2A, B**). We observed that cells generated inwardly directed contractile tractions against the substrate, even when the nucleus was highly deformed and transiting through the constriction. (**Figure 5A, B**). Cells on substrates with confinements were more polarized, displayed increased contractile forces, generated greater strain energy, had a larger maximum traction, and deformed their nucleus to a greater extent than cells on substrates without confinements (**Figure 5C - E, Supplemental Figure 2C**). When compared to cells in straight channels, we also observed that cells encountering constrictions exert more force along the Y axis (parallel to channel) and along Z but not X (perpendicular to the channel walls, **Figure 5F - H**).

**Figure 5.**
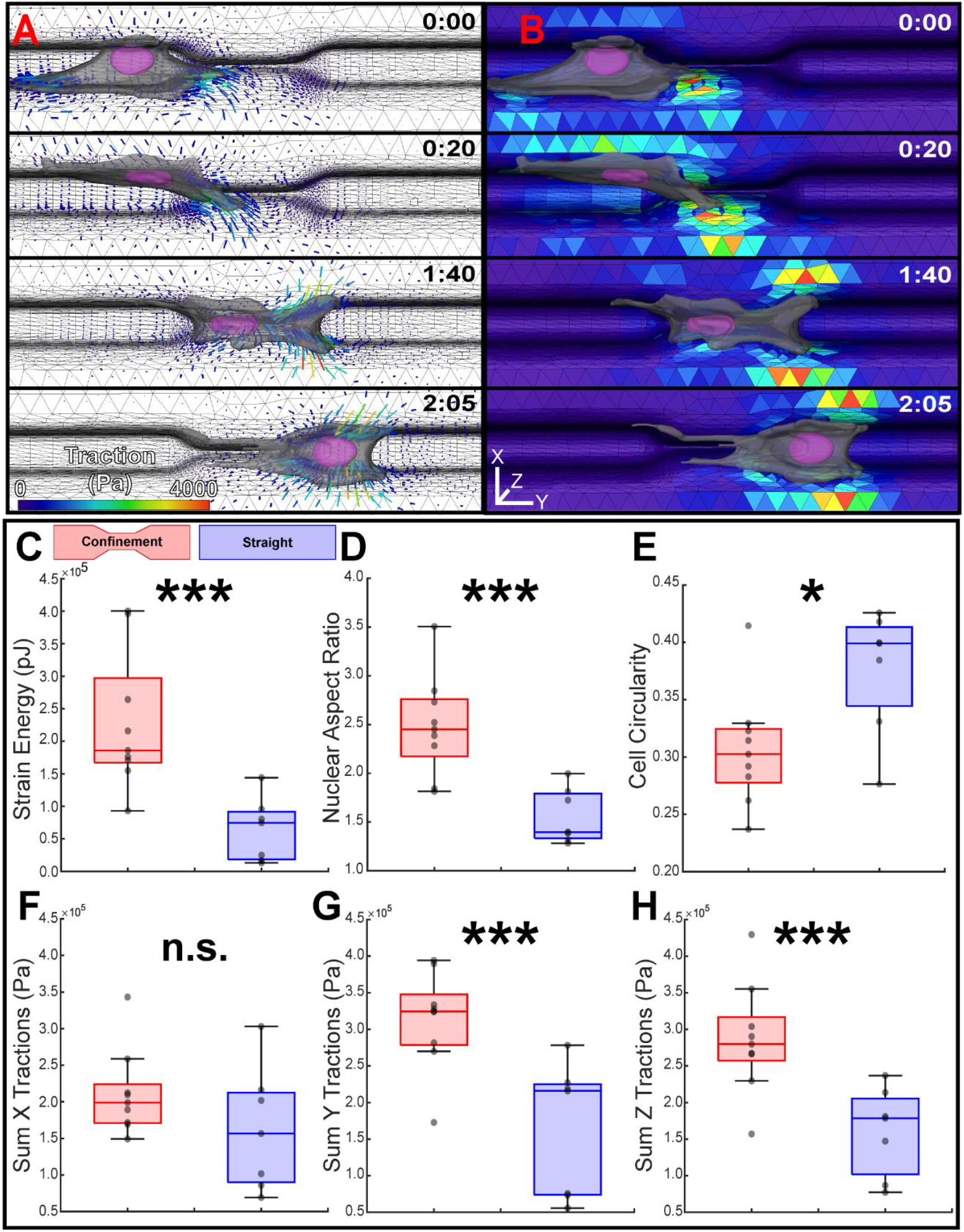
Measurement of cellular forces, nuclear deformation, and cell shape in straight channels and in channels with confinements. **A)** Montage images of a segmented cell (grey) and nucleus (magenta) migrating through a confinement with overlaid traction vectors. The magnitude of the tractions is color coded while the vector length is scaled for easier visualization. **B)** Contour map of the magnitudes of the traction vectors mapped to the element surfaces where they are computed. Time is hours:minutes. **C-E)** Average strain energy, 2D projected nuclear aspect ratio, and cell circularity for cells within compliant confinements (red) as opposed to straight channels without confinements (blue). **F-H)** Summed traction magnitudes in each direction during migration through compliant confinements and straight channels. All values are measured from the average of the measurements of each cell during migration through either a confinement or straight channel where each data point represents the average value from one cell (N = 9 for confinements, N = 7 for straight channels from two independent biological replicates). *** is p<0.001, ** is p<0.01, and * is p<0.05 as determined by Students T Test.

### The force to deform the nucleus during confined migration does not result from substrate induced reaction forces

Previous literature suggests that to deform the nucleus for transit of restrictive pores, the cell physically pulls or pushes the nucleus through the confinement (5–8). These mechanisms would require the nucleus to impart an outward deformation or pressure on the microchannel during transit which conflicts with our observations of strongly contractile deformations along the channel wall. To resolve this discrepancy, we measured the magnitude and direction of the traction forces along the channel walls within five microns of the nuclear periphery (**Figure 6A**). When averaged, we found the region immediately surrounding the nucleus is consistently under low magnitude negative force (and thus pulled towards the nucleus) throughout transit of the confinements and not significantly varied from cells in channels without confinements (**Figure 6B**). These observations are supported by similarly measuring the deformation of the substrate during transit where we observed no significant outward deformations (**Supplemental Figure 2D**), and by direct 3D visualization of the forces produced by the cell (**Figure 6C-E**). Thus, our observations are consistent with a model of confined migration whereas the force necessary to deform the nucleus arises from internal cytoplasmic components (**Figure 6F**) rather than pushing or pulling the nucleus through the confinement (**Figure 6G**).

**Figure 6.**
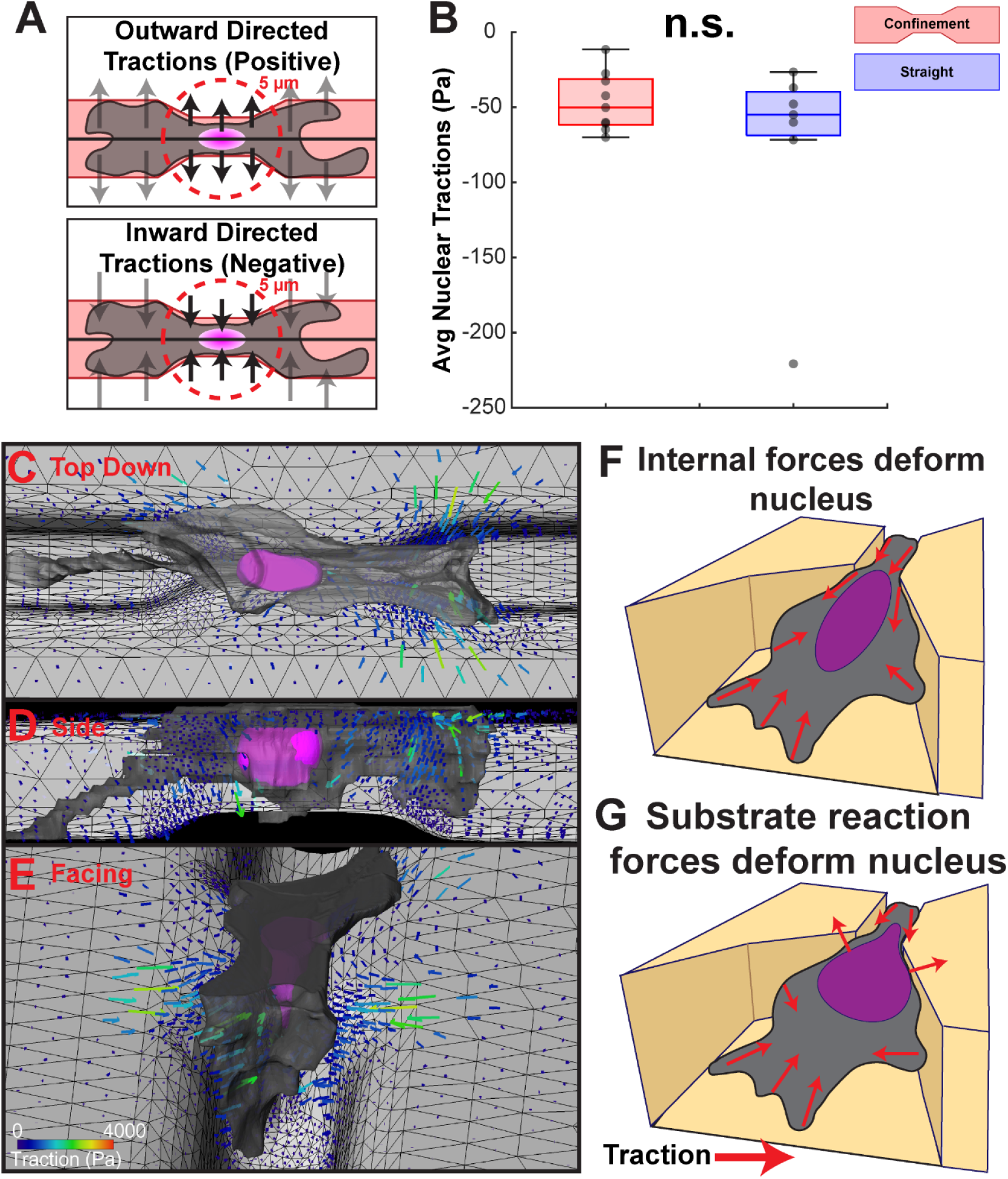
Cell-generated contractile forces rather than reactive substrate forces deform the nucleus when transiting through the constriction. **A)** Cartoon showing how the directionality of the inward and outward tractions along the axis of migration are determined. Forces pulling towards the center line of the channel are negative while those pushing outward are positive. **B)** Measurement of the average traction forces along the axis of migration within 5 microns of the nuclear surface for cells migrating through confinements (red) and straight (blue) channels. Average nuclear tractions were not significantly different as determined by Students T Test (N = 9 for confinements, N = 7 for straight channels from two independent biological replicates). **C-E)** Representative views (XY, YZ, and a tilted view respectively) of the exerted tractions when the nucleus is enclosed in the channel. **F, G)** Cartoons demonstrating of the expected force patterns when internal forces deform the nucleus (F) vs. when the nucleus is deformed by reaction forces from the substrate (G).

## Discussion

In this work, we generated micromolded substrates with sub-cellular-sized features and physiologically relevant stiffness. We used these substrates to measure the traction forces produced by cells as they migrate through restrictive confinements. In contrast to polymerized collagen (27, 30) or molded hydrogels (19, 20, 26), compliant PDMS is dimensionally stable and does not swell after polymerizing or in response to changes in temperature or osmolarity. Additionally, our sacrificial micromolding approach makes use of standard microfabrication reagents and commonly available sugars. Thus, a single silicon template can generate many low-cost substrates for future studies. We envision that these methods will be useful throughout the field of soft lithography both for biological studies and for material science.

By combining these substrates with high-resolution live-cell imaging and 3D TFM, we measured the forces exerted by cells as they transit through confinements. These measurements allowed us to assess the origin of the force necessary to deform the nucleus during transit. Previous studies have speculated that the nuclear deformation force could be derived from the reaction forces of the channel wall pushing against the cell as it pulls or pushes the nucleus (5–8) through the constriction. Alternatively, the cell may deform the nucleus via actin (3), microtubule (4), or intermediate filament networks (9). For the geometries and cell types studied here, we find that the nuclear deformation force does not arise from reaction forces within the confinement, but rather from cytoskeletal networks that also pull inward on the channel walls (**Figure 6**). Interestingly, cells migrating within the channels adopted a common “Y-shaped” protrusive structure that became compressed within the constriction and reformed upon exit (**Supplemental Movie 4**). Qualitatively, we also observed that the entire cell become more polarized once the leading edge encountered the confinement, suggesting that constriction at the leading edge triggers a global reorganization of the cytoskeleton to enable efficient transit. On 2D substrates, cell shape has been shown to regulate traction force generation (31), stem cell differentiation (32), and proliferation (33, 34). Given that 3D matrix geometry can similarly pattern cell shape and traction forces we anticipate that it may have an equally profound effect on cell phenotype. Future work to examine the relevance of these observations across cell types, ECM conditions, and other geometries will be critical to better understand how cell phenotype and migration strategies are influenced by both cell-intrinsic and cell-extrinsic factors such as substrate rigidity and geometry.

A key advancement in our methodology is the adaptation of our TFM pipeline to resolve cellular tractions on arbitrary 3D geometries (**Figure 3, Figure 4**). We envision this methodology will be broadly extensible to a variety of 3D systems and length scales and could be useful to address other mechanobiology questions related to the effects of substrate geometry (e.g. cell responses to positive vs. negative curvatures (35–37)) and stiffness. However, like other TFM methodologies, this approach has inherently low throughput and requires high resolution, 3D live cell microscopy to reproducibly measure the forces generated by cells. This makes it challenging to apply these methods to approaches such as high-throughput screening or to detect minor changes in traction force patterns across populations of cells (38, 39). Although we found that cells remained confined within the channels while migrating and similar open-top geometries have been used to study confined migration previously (26), future studies will be necessary to evaluate the effect of constrictions within fully enclosed 3D microchannels on the tractions exerted by cells. Lastly, PDMS elastomers are useful compliant substrates for micromolding as they faithfully preserve microchannel features despite osmolarity and temperature fluctuations however they are non-degradable by cellular processes such as proteases which have been implicated in the cellular response to confinement in more *in vivo* like collagen substrates or degradable hydrogels (28, 40). Our studies with fibronectin-coated microconfinements and 3D TFM have demonstrated the utility of these devices to study confined migration and to quantitatively measure the forces needed to deform the nucleus during cellular transit of non-directionally migrating mesenchymal cells. It is likely that other cell types (e.g. those undergoing amoeboid migration), those migrating in response to a directional signal, or those utilizing different adhesive ligand receptors may apply different force patterns while migrating through constrictions. The compliant PDMS micropatterned substrates introduced here will enable new microdevices that better control for substrate rigidity, adhesive conditions, and micron-scale 3D geometry to study these alternative propulsive strategies. Native tissues span a large range of rigidity, pore size, and adhesivity; each of which can be independently tuned in this system. Future applications will reveal how cells utilize traction forces to migrate through complex 3D environments.

## Materials and Methods

### Photolithography

Generation of silicon wafers with photoresist patterns was achieved using standard negative photolithography in a cleanroom as previously described (41). Briefly, a chrome on quartz mask was produced with our patterns of interest in negative photopatterning (Digidat, inc.). Then SU-8-2015 (Kayaku Advanced Materials) photoresist was deposited on 4-inch silicon wafers (University Wafer 452) by dispensing 4 mL of resist before spinning the wafer on a spin coater at 1000 RPM for 30 seconds to achieve a film thickness of approximately 30-microns. The wafer coated with photoresist was baked at 95°C for 5 minutes on a hotplate. Next, the wafer was transferred to a Karl Suss MA6/BA6 Mask Aligner equipped with a 350 nm longpass filter and aligned with the chrome mask with desired patterns before exposing to 150 mJ/cm^2^ UV light. Once complete, the wafer was post exposure baked for five minutes at 95°C. Finally, the wafer was developed using SU-8 developer on a shaker table for 5 minutes before rinsing with iso-propyl alcohol and drying with pressurized nitrogen. Before use, the silicon wafer was exposed to trichlorosilane (Sigma-Aldrich 448931) by adding a drop to a glass slide placed in a vacuum chamber and left overnight within a chemical fume hood. The silane vaporizes and deposits a layer of silane on the wafer which facilitates removal of the silicon wafer from later PDMS molds by rendering the surface nonadherent.

### Sacrificial Micromolding

Compliant microchannels were generated by first fabricating a rigid PDMS cast of the silicon wafer containing the desired micropatterns. Sylgard 184 (Dow 1317318) was prepared at 10:1 (Part A to Part B by weight) and mixed thoroughly before degassing for 15 minutes in a tabletop vacuum chamber. After degassing, Sylgard 184 was added to the silicon wafer that was previously placed in a 10cm plastic petri dish. Approximately 50 mL of degassed Sylgard 184 was added to the wafer, which is pushed to the bottom of the petri dish to ensure it is fully submerged. The entire dish was placed in a tabletop vacuum chamber for 30 minutes to remove any air bubbles before placing in a 60°C oven overnight. The following day, after fully curing, the petri dish was removed from the PDMS and silicon mold by carefully snipping the petri dish with wire cutters while taking care not to crack the silicon wafer (for reuse and generation of subsequent molds). Once extracted, the PDMS and silicon molds were separated, the silicon mold was cleaned with compressed nitrogen gas and stored in a covered plastic petri dish, while the PDMS replicate is treated with trichlorosilane overnight as described above. After silane treatment, the PDMS replicate is ready for use in producing sugar molds.

Sugar molds were prepared using a molten mixture of granulated sugar and light corn syrup (Karo). Briefly, 30 grams of granulated sugar was added to a clean and completely dried 400 mL glass beaker. 15 grams of light corn syrup was added and mixed into the sugar using a metal spatula. The sugar mixture was then microwaved for a total of 1 minute and 45 seconds or until the solution turns a dark amber, is bubbling, and smells of caramel. The molten solution was then immediately poured on the PDMS replicate mold surrounded by a silicon rubber gasket punched from a silicon rubber sheet. A metal spatula was used to gently scrape the rigid PDMS micropattern to ensure the molten sugar settles into the pattern and to remove any bubbles. Finally, a 25 mm coverslip was added to the top of the molten sugar mixture and pressed down onto the silicon gasket to make a flat top. The sugar mold was left to cool for one hour before demolding from the PDMS template.

To generate compliant microchannels, a solution of compliant PDMS was prepared by mixing rigid Sylgard 184 and compliant Sylgard 527 (Dow 1696742). Briefly, Sylgard 184 was prepared at 10:1 (Part A to Part B by weight) and mixed thoroughly before degassing for 15 minutes in a tabletop vacuum chamber. While degassing, a Sylgard 527 solution was prepared at 1:1 (Part A to Part B by weight), mixed thoroughly and degassed for 15 minutes in a tabletop vacuum chamber. Once both solutions are degassed, they were mixed in a 20:1 ratio (Sylgard 527 to Sylgard 184) by weight and mixed thoroughly before degassing for 15 minutes in a tabletop vacuum chamber. Ultra-compliant PDMS (∼2 kPa) was generated by using solely Sylgard 527 solution. Once degassed, approximately 25 µL of the compliant PDMS mixture was immediately added to the demolded sugar template. A 50-micron thick circular metal shim was added on top of the sugar mold and a plasma cleaned #1.5 25 mm coverslip was added to complete the stack. The stack is inverted and then carefully set on a parafilm sheet before 50 grams of weight is added to each mold and allowed to cure at room temperature for at least 48 hours. Once cured, the mold was immersed in a water bath and allowed to shake for 10 minutes before the top glass coverslip w<u>as</u> removed. A further 15 minutes of immersion was followed by carefully removing the silicon gasket using a razor blade and then the metal shim by gently peeling it from the 25 mm coverslip. The remaining coverslip and attached sugar mold were then immersed in a water bath for a further 1 hour or until the sugar is completely dissolved. Once dissolved, the coverslip with micropatterend PDMS mold was removed, washed several times with clean diH20 and stored in diH20 in a 6 well plastic dish before use. It is important after this point to ensure that the molds are not allowed to dry out as capillary forces from evaporating liquid can collapse the channels.

### Addition of fluorescent beads

To add fluorescent beads to the substrates, we removed the diH20 and washed the surface with 100% EtOH. We then treated the surface with 2 mL of a 10% by volume APTES (Thermo-Scientific 430941000) in 100% EtOH for 30 minutes on a shaker table. The APTES solution was removed from the substrates, and they were washed 3x with 100% EtOH before 2 mL of a 1/1000 solution of carboxylate polystyrene fluorescent beads (Thermo-Fisher F8801) and 1 mg/mL EDC (Thermo-Fisher E7750) in diH20 was added to the substrates, which were then allowed to rock on a shaker table for 20 minutes. The bead and EDC solution was then removed, and the surface was washed with diH20.

### Addition of fibronectin and sterilization

To add fibronectin and sterilize the substrates, we placed the substrates upside down into a fresh, sterile plastic 6-well dish containing 1 mL of 10 µg/mL of fibronectin (Gibco 33010018) inside of a tissue culture hood. The plastic 6-well dish was then irradiated with UV light for 30 minutes to sterilize the substrates while the fibronectin coated the surface. Once irradiated, the substrates were washed with sterile PBS 3x times.

### Plating cells

Substrates were placed into a 35 mm metal imaging dish and 1 mL of culture media containing approximately 5000 cells and 1 µM SiR-Hoescht (Spirochrome) was added before a sterile plastic lid from a 35 mm plastic cell culture dish was used to cover the dish. The imaging dish was then fitted into a custom PDMS mold attached to a centrifuge bucket, and the cells were spun down into the microchannels at 48 RCF for 5 minutes before the dish was rotated 180 degrees and spun for an additional 5 minutes at 48 RCF. Cells were allowed to attach for 2 hours before imaging.

### Microscopy

Imaging experiments were conducted using different microscopes depending on the necessary resolution, optical sectioning, and throughput. All microscopes were equipped with a live-cell imaging chamber maintained at 37°C and 5% CO_2_.

Low resolution imaging was performed on a Nikon TI2 inverted widefield microscope with a 10x 0.45 NA Plan Apochromat air objective. Cells were imaged every 5 minutes for 18-24 hours under brightfield (100 ms exposure, 5% condenser lamp power) and Cy5 (100 ms exposures and 5% laser power) illumination. 15 fields of view were selected from each of four imaging dishes simultaneously imaged.

Point-Scanning confocal imaging was performed on a Zeiss Airyscan 800 microscope using a 40x NA 1.3 EC Plan-Neofluar oil objective. The pinhole was set to 1 Airy-Unit, and pixel dwell time and laser exposure were set to maximize signal dynamic range.

63x Spinning disk confocal microscopy was performed with a Leica Dragonfly microscope at the Microscope Services Laboratory at UNC Chapel Hill. The microscope was equipped with an iXon Life 888 EMCCD camera and a 63x/NA 1.4 HC PL APO oil objective. The pinhole was 40 microns. 15-20 fields of view were selected on each imaging dish and 45-micron 3D volumetric stacks with a z step of 800 nm were collected at each position, every 5 minutes. Three color channels were collected sequentially: GFP (Lifeact-GFP) 2.5% laser power exposure 50 ms exposure, RFP (Fiducial Beads) 5% laser power 25 ms exposure, and Cy5 (DNA) 5% laser power 25 ms exposure. Following traction force experiments, cells are removed from the substrate by adding 100 µL of 0.1% b/w sodium dodecyl sulfate (Fisher 28364).

### Cell Culture and Reagents

All cells were cultured and imaged in high-glucose DMEM (Gibco 11965092) supplemented with 10% FBS (MedSupply Partners 62-1300-1) and 100 U/mL of Penicillin/Streptomycin (Gibco 15140122) at 37°C and 5% CO2. Cells were split and passaged using 0.25% trypsin-EDTA (Gibco 25200056).

IA32 Lifeact-GFP cells were generated as described previously (42). IA32 cells were grown to confluence before FACS sorting to enrich the population for cells with a moderate expression of Lifeact-GFP.

A mouse embryonic fibroblast (MEF) clonal line (JR20) was established from a previously described ARPC2 conditional knockout mice (43). Lentivirus for the stable expression of cytoplasmic GFP was generated through the transfection of pCMV-V-SVG, pRSV-REV, pMDLg/pRRE, and pLL5.0-eGFP (500 ng each) plasmids into HEK293FT cells using X-tremeGENE HP transfection reagent (Roche). Lentivirus was harvested 72 hours later and subsequently used to infect JR20 MEFs supplemented with 4 μg/mL of Polybrene (Santa Cruz). Roughly 72 hours following the addition of lentivirus, JR20s that underwent transduction and were expressing cytoplasmic GFP were isolated via FACS.

### Indentation Rheology

Measurement of the elastic modulus of the PDMS substrates was performed using indentation rheology with tungsten carbide microspheres (44). Briefly, 300-micron tungsten microspheres (NEMB) were carefully dropped on the surface of a flat PDMS substrate coated with 100-nm fluorescent microspheres. The indentation of the tungsten microsphere was imaged using a confocal microscope and the maximum deformation of the substrate was measured in Fiji by viewing a projection of the XZ axis. Then the height of the PDMS substrate was calculated by measuring the height of the surface of the PDMS substrate and the minor imperfections in the PDMS at the bottom of the glass coverslip. These values were inputt into the workflow as described in (44) to obtain the elastic modulus of the substrate. We assumed a Poisson ratio of 0.5 for a linear elastic solid.

### Shear Rheology

Measurement of the loss and storage moduli of the PDMS substrates was performed using parallel plate rheology (45). Briefly, fully cured PDMS cylinders 8mm in diameter were prepared from larger disks and then transferred to a TA Instruments Discovery HR-3 hybrid rheometer. A strain sweep was performed at 1 rad/s to determine the linear viscoelastic regime. Subsequently, frequency sweep experiments were performed at 1.0% strain (within the linear viscoelastic regime) at an axial force of 0.1 N at 22 °C from 0.01 to 1000 rad/s. Storage and loss modulus were recorded at each measured frequency. Data above 100 rad/s was excluded as the substrates frequently broke and/or lost contact with the rheometer. The average moduli values at 1 rad/s are reported.

### Measurement of Cell Inducted Traction Forces in Complex Geometries

To measure the displacement field produced by cells deforming the substrate surface, we localized the 3D position of the fluorescent fiducial markers in our 3D volumetric imaging data and then determined the corresponding location of the bead in the cell free state using feature vector linking as described previously (11, 14). Briefly, the 3D volumetric stacks were loaded into a custom MATLAB script where a maximum likelihood estimator (MLE) fitting algorithm localized the 3D center of each fluorescent fiducial bead. To compute a displacement field for a given time frame, each deformed frame bead must be matched to its corresponding bead in the relaxed state. To link the fiducial beads, each deformed frame was compared to the reference frame (where the cell is removed via detergent) and the localizations were matched using feature vectors. An initial set of feature vectors were determined to compute the overall drift. Because addition of the detergent SDS causes a small non-uniform swelling of the substrate geometry, the swelling profile was determined by capturing the displacement of beads in frames where the cells were not present in the field of view. At least three of these swelling fields were averaged and then added to the localizations in the reference state. The resulting corrected “swelling-free” reference state was then used for subsequent frames to match fiducial beads. A second set of feature vectors were then determined to match the deformed and relaxed states. The resulting vector field is the displacement of the fiducial beads resulting from the cell induced tractions.

To compute the traction field from a given displacement field, finite element modeling was used to measure the mechanical response of a given geometry to point forces. Briefly, the 3D localizations of the fidiciual beads were used to generate a model of the surface with tetrahedral elements which was then converted into a STL using a custom MATLAB script. The STL was then loaded into Hypermesh (Altair Engineering) which was then used to smooth the mesh. Boundary conditions were added to the edges of the substrate preventing the outermost elements from deforming under loads. Finally, a custom TCL script was used to generate a subcase containing a single X, Y, or Z load on each triangular element face on the outer face of the model. The resulting set of load conditions is then solved using Optistruct (Altair Engineering).

We used a custom MATLAB script to convert the finite element solution into a Green’s matrix for solving the corresponding tractions from experimental displacement data. First the finite element solution for each load case was loaded into MATLAB and then converted into a matrix array holding the resulting displacements for a given unit load on a single element as each row. The matrix underwent singular value decomposition and was combined with the displacement field using Tikhonov regularization which outputs the traction field given a regularization parameter. The regularization parameter defines a tradeoff between the resulting traction field resolution and minimizing the residual between the forward solution and the experimental displacements. A value for the regularization parameter was determined by comparing the residual error of the experimental displacement field and the resulting forward displacement field predicted by the traction field such that the average residual error was equivalent to the 3D localization precision of our fluorescent beads. This regularization value was then used for all TFM datasets in the manuscript.

### Analysis and statistics

Analysis and statistics were performed using MATLAB and FIJI and described below. Transit time was measured as the time when the centroid of the nucleus first passes within 20 microns of the confinement center and the last time point it passes the 20 microns on the opposite side of the confinement center. Constriction width change was measured as the percentage width change of confinement walls from before the cell contacted the confinement, and the width change when the nucleus is closest to the center of the confinement. Strain energy was computed as previously described (12). Nuclear aspect ratio was computed as the ratio of the major to minor axis of a fit ellipse of the masked image of a nucleus. Cell circularity was computed as the scaled measure of the area to perimeter ratio which ranges from one for a perfect circular projection to a value of zero for a highly extended shape and is thus a proxy for cell polarization. The sum of the traction projections was computed by summing the magnitudes of the traction vector components. The average nuclear traction was computed by averaging the magnitudes of the X component of the tractions within 5 microns of the nuclear mask. The sign of the magnitude was flipped for values whose position was negative relative to the center of the channel. Thus, all positive values represent forces “pushing” outwards while negative values represent forces directed towards the center of the confinement. Average nuclear displacements were computed following the same scheme for the displacements within five microns of the nucleus. Maximum traction was computed as the maximum traction value magnitude reported for all elements at a given time point. Significance values for pairwise comparisons were performed using a two-sided Student’s T Test with Bonferroni multiple hypothesis correction where appropriate.

## Code and data availability

All code and data produced for this manuscript is available upon request.

## Acknowledgments

We thank Adam Palmer and all members of the Legant and Bear labs for helpful discussion and critical feedback.

This work was funded in part by grants from the National Institutes of Health (1DP2GM136653) awarded to W.R.L., (R35GM130312) awarded to J.E.B., and (1R35GM142666-01) awarded to F.A.L. and used to support J.L.R.. W.R.L acknowledges additional support from the Packard Fellowship in Science and Engineering.

The Microscopy Services Laboratory, Department of Pathology and Laboratory Medicine, is supported in part by P30 CA016086 Cancer Center Core Support Grant to the UNC Lineberger Comprehensive Cancer Center. The Andor Dragonfly microscope was funded with support from National Institutes of Health grant S10OD030223. The UNC Flow Cytometry Core Facility (RRID:SCR_019170) is supported in part by P30 CA016086 Cancer Center Core Support Grant to the UNC Lineberger Comprehensive Cancer Center. Research reported in this publication was supported in part by the North Carolina Biotech Center Institutional Support Grant 2012-IDG-1006. This work was performed in part at the Chapel Hill Analytical and Nanofabrication Laboratory, CHANL, a member of the North Carolina Research Triangle Nanotechnology Network, RTNN, which is supported by the National Science Foundation, Grant ECCS-1542015, as part of the National Nanotechnology Coordinated Infrastructure, NNCI.

