## Supplemental Information for "Measurement of cellular traction forces during confined migration"

### Supporting Information for
Measurement of cellular traction forces during confined migration

Max A. Hockenberry, Andrew J. Ulmer, Johann L. Rapp, Frank A. Leibfarth, James E. Bear*, Wesley R. Legant*

*Corresponding Authors: Wesley R. Legant, James E. Bear

**This PDF file includes:**

Supporting text
Figures S1 to S2
Legends for Movies S1 to S4

**Other supporting materials for this manuscript include the following:**

Movies S1 to S4

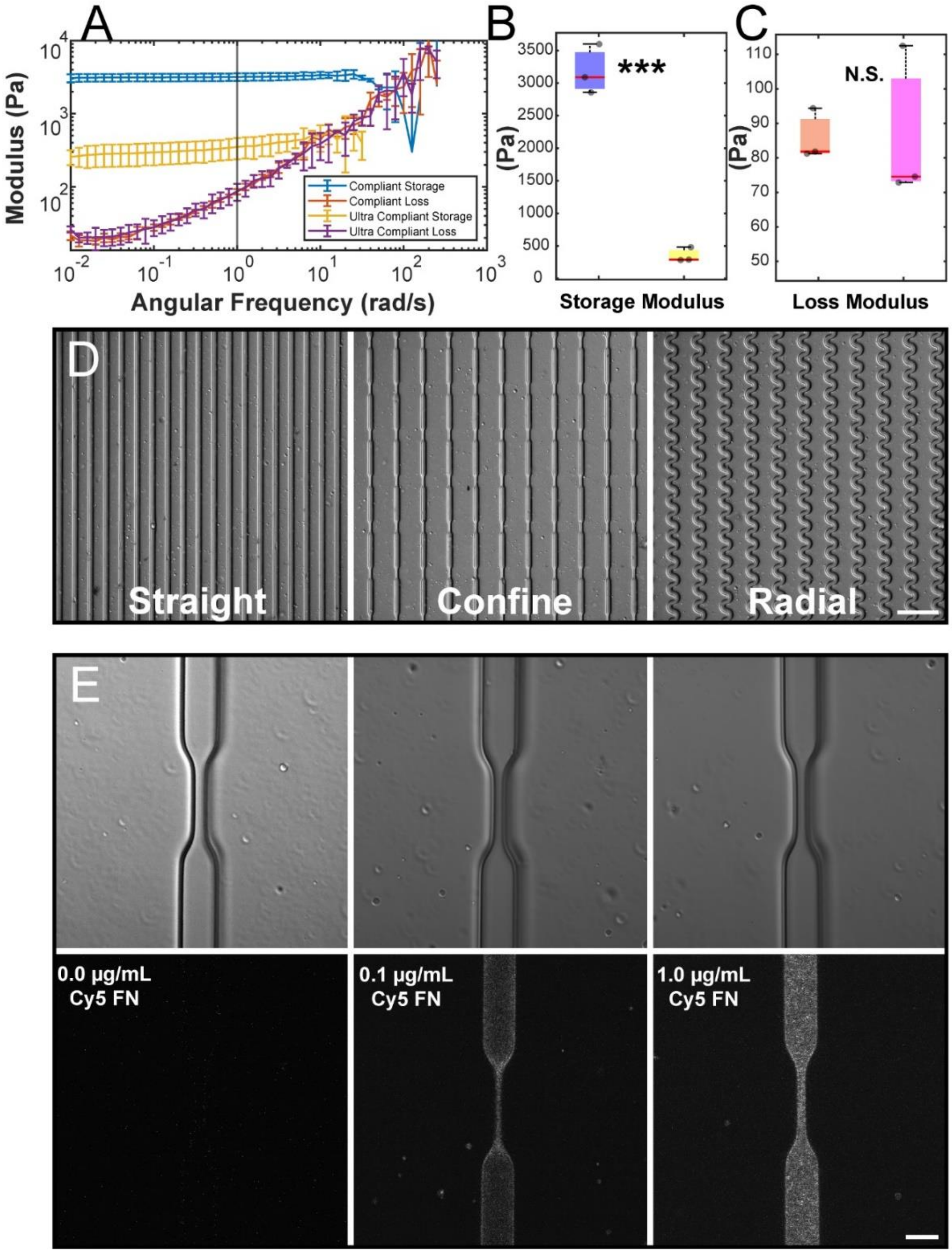


**Figure S1. Characterization of microchannel structures, ECM ligand attachment, and shear rheology**. **A)** Average storage and loss moduli as a function of angular frequency measured during three independent shear rheology experiments. **B, C)** Average storage and loss modulus values at 1 rad/s angular frequency for compliant and ultra-compliant rigidity substrates. **D)** Large field of view brightfield images of micropatterned substrates showing repeated geometric features. **E)** Confocal images of microconfinements with varying concentrations of Cy5 fibronectin attached to the surface. Increasing concentration of Cy5 FN results in increasing density of fluorescent signal. *** is $p<0.001$ as determined by Students T Test from three independent experimental replicates.

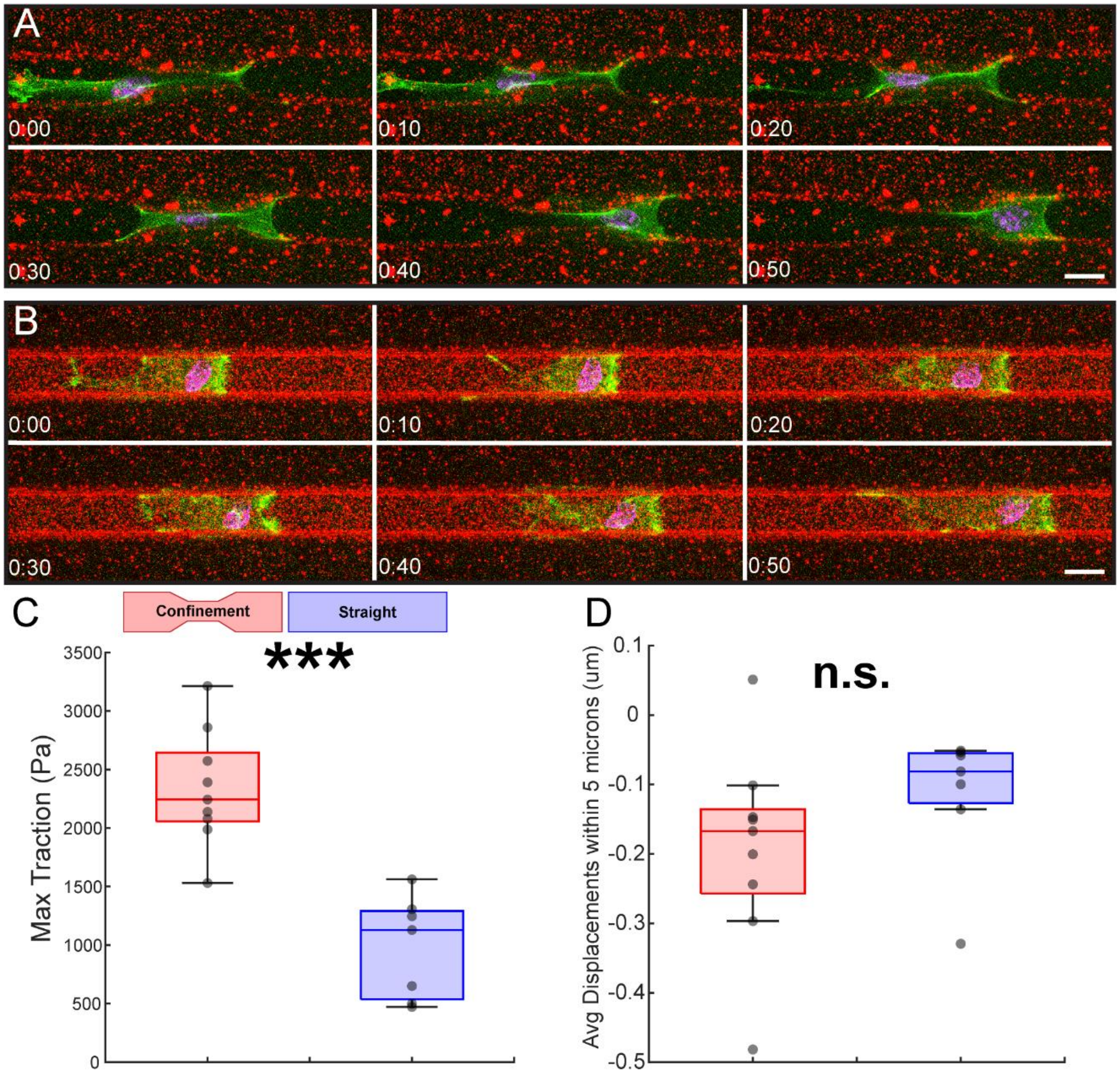


**Figure S2. Imaging of cell migration on compliant substrates with and without constrictions. A, B)** 3D volumetric images projections of IA32 LifeAct-GFP (green) expressing cells labeled with SiR-DNA (magenta) migrating through constricting (A) or straight (B) channels. Scale bar = 20 microns and the time is hours:minutes. **C)** shows the time-averaged maximum reported traction for cells migrating in confinements (red) and straight (blue) channels. **D)** shows the measurement of the average displacement along the axis of migration within 5 microns of the nuclear membrane for cells migrating through confinements (red) and straight (blue).Negative values are directed towards the nuclear membrane while positive values are away from the nucleus. All values are measured from the average of the measurements of each cell during migration through either a confinement or straight channel where each data point represents the average value from one cell (N = 9 for confinements, N = 7 for straight channels from two experimental replicates). *** is P<0.001, ** is P<0.01, and * is P<0.05 as determined by Students T Test.

**Movie S1 (separate file). Low magnification widefield imaging of constriction transit events**. Movie shows a single field of view captured during overnight live cell migration experiments with individual transit events tracked and labeled with colored dots and lines. On average, 60 fields can be captured across four separate substrates. Cells are IA32 LifeAct-GFP (not shown) MEFs stained with SiR-DNA (red). Brightfield imaging is use to visualize both the cells and channel structures. The field is captured every five minutes. Scale bar = 200 microns.

**Movie S2 (separate file). Representative images of cells in rigid and compliant microchannels transiting through constricting regions.** Cells experiencing compliant constrictions transit slower, deform the substrate significantly, and have a higher nuclear aspect ratio. The movie is of two representative regions of interest on a compliant and rigid substrate with 5-micron wide, 40-micron long confinements imaged via low magnification. Cells are IA32 LifeAct-GFP (not shown) MEFs stained with SiR-DNA (red). Brightfield imaging is use to visualize both the cells and channel structures. The field is captured every five minutes. Scale bar = 20 microns.

**Movie S3 (separate file). Measurement of cellular traction forces exerted by a cell migrating through a microconfinement.** The movie shows two views of a 3D volumetric dataset collected at 63x magnification on a spinning disk confocal microscope. The top two panels are the XY and YZ projections of the raw data of a JR20 MEF labeled with a cytoplasmic GFP (green) and SiR-DNA (orange) migrating on a substrate labeled with 100-nm fiducial fluorescent beads (red). As the cell migrates, it deforms the substrate causing the beads to move. The nucleus is deformed during transit and upon exit the cell continues migrating away from the confinement. The bottom two frames are the same raw data but overlaid with the traction vectors and STL mesh of the substrate geometry. Traction magnitudes are color coded and the vectors lengths are scaled for visualization. Frames are captured every five minutes. Scale bar = 20 microns.

**Movie S4 (separate file). Cell migrating through narrow confinement deforms the nucleus and generates inwardly directed traction forces.** The movie shows a 3D volumetric dataset collected at 63x magnification on a spinning disk confocal microscope. The movie shows an IA32 LifeAct-GFP MEF stained with SiR-DNA which is segmented for the cell (grey) and nucleus (magenta). The cell migrates through a confinement with the geometry shown as a STL mesh (white). The deformation of the cell’s nucleus prior to nuclear confinement by the channel is highlighted. During transit of the confinement, the tractions immediately around the nucleus are highlighted to demonstrate their inward direction. The tractions are shown as vectors with their magnitude color coded and vector lengths scaled for visualization. Frames are captured every five minutes.
